# UNKAI: A protein functional identity prediction model based on ESM-C latent representations and the attention mechanism

**DOI:** 10.64898/2026.05.02.722384

**Authors:** Kotaro Ukai, Suguru Fujita, Tohru Terada

## Abstract

The rapid advancement of genome sequencing technologies has led to the accumulation of a vast number of protein sequences in public databases. However, a significant proportion of these proteins remain functionally uncharacterized. Concurrently, the expansion of protein sequence data has enabled the development of protein language models (pLMs). By distilling billions of years of evolutionary history into a latent representational space, these models have acquired an unprecedented capacity to predict both the tertiary structures and functions of proteins. In this study, we developed a deep learning-based method to predict whether two proteins catalyze the same enzymatic reaction. Our approach leverages latent representations generated by ESM Cambrian (ESM C), a state-of-the-art pLM, which are then processed through a neural network architecture integrating an attention mechanism. Our method outperformed existing approaches, including those based solely on full-length sequence similarity. Notably, it also surpassed our previous LightGBM-based model, which relied on structural similarity scores derived from AlphaFold-predicted models. Analysis of the attention weights reveals that our model autonomously highlights biologically significant sites, such as catalytic and binding residues. This demonstrates that integrating pLMs with attention mechanisms can enhance the accuracy and interpretability of protein function prediction while eliminating the need for manual feature engineering.

## 1. Introduction

Owing to the rapid advancement of genome sequencing technologies, public databases— such as the Reference Sequence (RefSeq) collection of the National Center for Biotechnology Information (NCBI)—now contain hundreds of millions of protein sequences [1]. However, fewer than 0.2% of these proteins have been experimentally characterized [2]. To bridge this gap and facilitate the discovery of proteins with novel functions, efficient computational methods for protein function prediction are essential.

Traditionally, protein function prediction has relied on sequence alignment tools, such as BLAST, to identify homologous proteins under the premise that the sequence similarity implies functional similarity [3]. While many recent computational methods frame function prediction as a multi-class classification problem [4–7], we focus on a pairwise approach inspired by these alignment-based methods to determine whether two enzymes catalyze the same reaction involving the same substrate. We previously implemented this pairwise approach using a classical machine-learning-based model referred to as “FUnctional identity of protein prediction by JoIning Sequence ANd structure feature (FUJISAN)”[8]. This model integrated full-length sequence similarity scores with structural similarity scores derived from the domain-level comparison of structural models predicted by AlphaFold2 [8,9]. FUJISAN demonstrated superior performance not only over a sequence identity-based baseline but also over models utilizing deep-learning-based embeddings from DeepFRI [10] and ESM-2 [5]. Despite its high performance, FUJISAN’s reliance on structure-based features limits its scalability to massive datasets due to the substantial computational overhead associated with structure prediction, domain decomposition, and pocket detection. However, such limitations are inherent to a “hypothesis driven feature engineering” approach, where specific biological features must be manually defined and computed.

To overcome these limitations, we propose a novel deep learning framework that enables end-to-end prediction by integrating latent representations from ESM Cambrian (ESM C) [11], a state-of-the-art protein language model (pLM), with an attention pooling mechanism [12,13]. This mechanism enables the model to prioritize functionally critical residues during the prediction process. By employing a Siamese network architecture [14] with a learnable distance function, our method captures complex, non-linear relationships between protein pairs without the need for explicit structural modeling or alignment. This study demonstrates that our end-to-end approach not only streamlines the computational pipeline but also achieves superior accuracy in predicting protein functional identity.

## 2. Materials and methods

### 2.1. Dataset

In this study, protein function is defined by the Rhea ID, which is uniquely associated with a specific pair of substrate and product [15]. Consequently, a protein pair with identical function possesses the same Rhea ID, indicating that both proteins catalyze the same reaction involving the same chemical transformation. Following the methodology of our previous study [8], we retrieved 232,692 sequences annotated with Rhea IDs from the SwissProt database [2]. An all-against-all BLAST search was performed on this sequence set and all hits with E-values less than 10 were retrieved. From these, only pairs consisting of proteins with up to 2,048 residues were retained for further analysis to accommodate the sequence length constraint of ESM C. Subsequently, these protein pairs were categorized into two groups: positive pairs, consisting of proteins sharing the same Rhea ID, and negative pairs, consisting of proteins with distinct Rhea IDs. From each group, 100,000 pairs were randomly sampled to construct the final dataset.

### 2.2. Model architecture

Adopting a Siamese network architecture [14], the proposed model comprises three main components: an embedding layer, an attention pooling layer, and a multi-layer perceptron (MLP) classifier (Figure 1A). In the embedding layer, the pre-trained pLM, ESM C [11], is employed to generate a latent representation of the input sequence with dimensions of *D* × *L*. Here, *D* and *L* denote the fixed dimensionality of the latent representation (*D* = 2,560) and the sequence length, respectively. Subsequently, the variable-length latent representation **X** ∈ ℝ^*D*×*L*^ is aggregated into a fixed-size vector **v** ∈ ℝ^*D*^ within the attention pooling layer, as illustrated in Figure 1B. Let **x**_*j*_ ∈ ℝ^*D*^ be the *j*-th column vector of **X** corresponding to the embedding of residue *j* . For each **x**_*j*_, an attention logit *e*_*j*_ is computed via a linear transformation:

**Figure 1.**
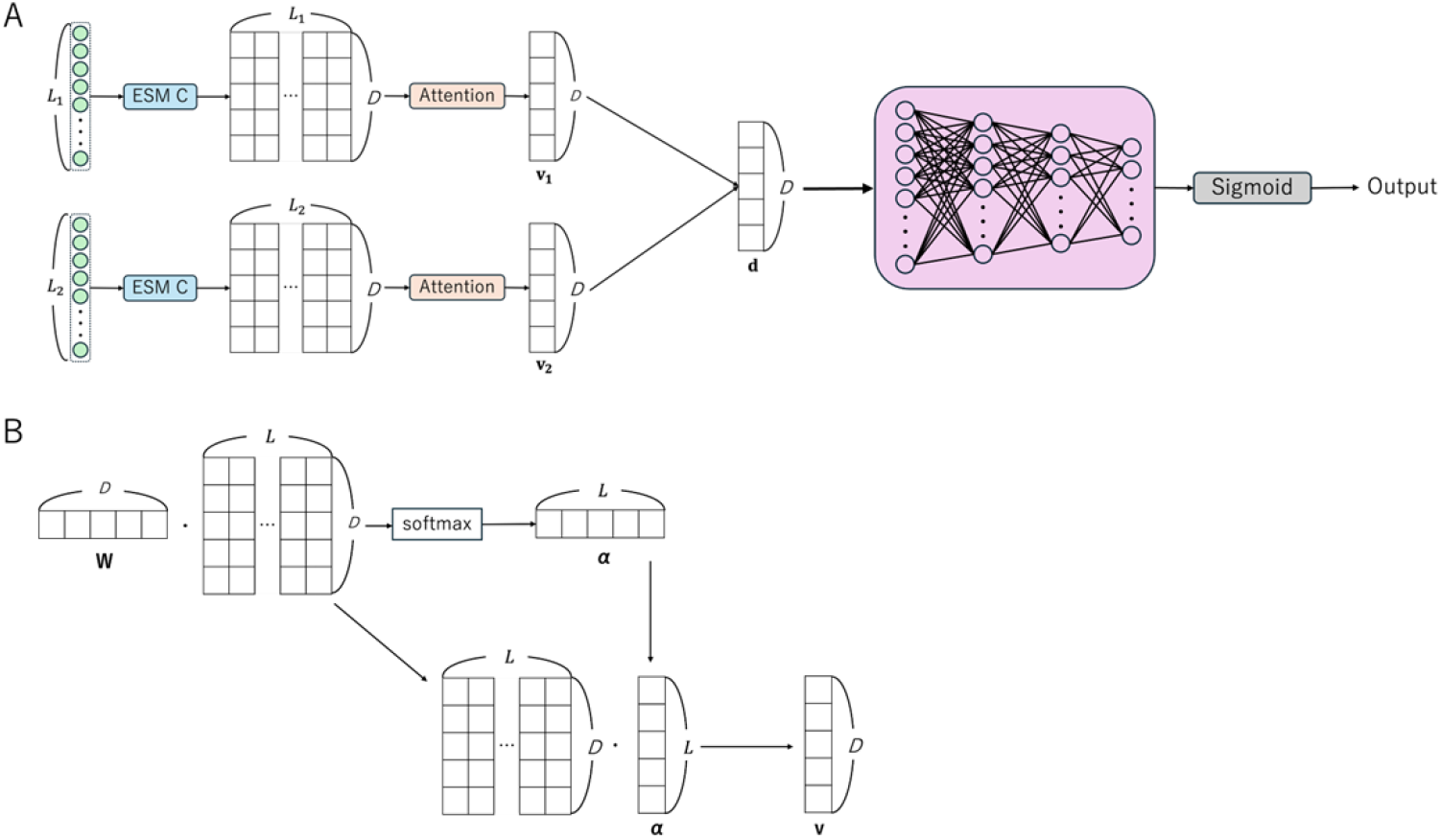
Model architecture. (A) Overall architecture of the proposed model, UNKAI, comprising an embedding layer, an attention pooling layer, and a multi-layer perceptron (MLP) classifier. In the embedding layer, amino acid sequences of the input protein pair (proteins 1 and 2) are converted to ESM-C latent representations with dimensions of *D* × *L*_1_ and *D* × *L*_2_, where *D* = 2,560 and *L*_1_ and *L*_2_ denote the sequence lengths. These representations are aggregated into fixed-size vectors **v**_1_ and **v**_2_ within the attention layer. The element-wise absolute difference **d** (where *d*_*k*_ = |*v*_1,*k*_ − *v*_2,*k*_|) is then calculated and fed into the MLP classifier. The final MLP output is passed through a sigmoid function to obtain the probability of functional identity. (B) Schematic representation of the computation within the attention pooling layer.

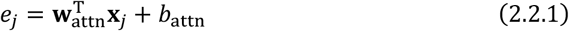

Here, **w**_attn_ ∈ ℝ^*D*^and *b*_attn_ are learnable parameters. The attention weights *α*_*j*_ are obtained by normalizing the logits *e*_*j*_ using the softmax function:

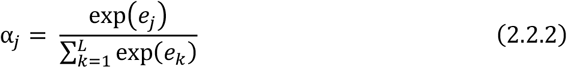

Finally, the aggregated feature vector **v** is computed as the weighted sum of the residue embeddings:

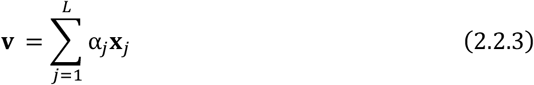

This process is applied independently to each protein of the input pair using shared parameters, yielding two representative vectors **v**_1_ and **v**_2_, corresponding to input 1 and input 2, respectively. To capture the relationship between the two proteins, the element-wise absolute difference **d** ∈ ℝ^*D*^ between **v**_1_ and **v**_2_ is calculated and fed into the MLP classifier:

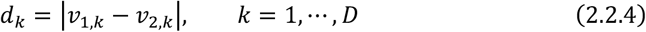

where *v*_1,*k*_ and *v*_2,*k*_ denote the *k* -th elements of **v**_1_ and **v**_2_, respectively. The MLP classifier consists of four fully connected layers. The input dimension is 2,560, matching the dimensionality of **d**, followed by three hidden layers with 1,599, 781, and 117 units, respectively. Each layer is followed by batch normalization, a ReLU activation, and a dropout layer with a rate of 0.302766. The final MLP output is passed through a sigmoid function to obtain the probability of functional identity, constrained between 0 and 1.

### 2.3. Model training and hyperparameter optimization

The positive (identical function) and negative (distinct function) pair datasets were independently split into training (70%), validation (15%), and test (15%) subsets. Subsequently, the corresponding subsets from both groups were combined and randomly shuffled to form the final training, validation, and test sets. This procedure ensured that each set contained an equal number of positive and negative pairs, providing a balanced distribution for both model training and evaluation. The model was trained using the Adam optimizer [16] with a learning rate of 3.33 × 10^−4^ and the binary cross-entropy (BCE) loss function, defined as follows:

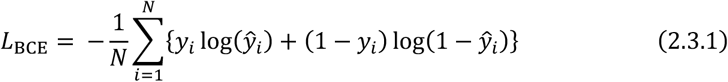

where *N* denotes the number of samples in a batch, *y*_*i*_ ∈ {0,1} is the ground-truth label, and ŭ_*i*_ represents the predicted probability of functional identity for the *i*-th pair. Training was performed for 10 epochs with a batch size of 64. Within each batch, sequences were padded to a uniform length, and an attention mask was applied to ensure that padded positions did not contribute to the attention weights or the resulting aggregated representations. The model achieving the highest F1 score (see below) on the validation set was saved.

To maximize the model’s predictive performance, hyperparameter optimization was performed via Bayesian optimization using the Optuna framework [17]. The search space encompassed the number of units in the three hidden layers (*n*_1_, *n*_2_, *n*_3_), the dropout rate, and the learning rate. We explored the following ranges for hyperparameter optimization: hidden layer sizes *n*_1_ ∈ [512,2048], *n*_2_ ∈ [256, *n*_1_], and *n*_3_ ∈ [64, *n*_2_] dropout rate between 0.1 and 0.5 and learning rate from 10^−5^ to 10^−2^, the latter of which was sampled log-uniformly. A total of 50 optimization trials were conducted, with the objective of maximizing the F1 score on the validation set. To improve computational efficiency, we employed a Median pruner strategy to early-terminate unpromising trials. The final model was trained using the optimal hyperparameter configuration identified during this process and subsequently evaluated on the independent test set. The resulting model is named UNKAI (protein fUNKtionAl Identity prediction). In Japanese, *unkai* refers to a “sea of clouds,” a breathtaking phenomenon often viewed from the summit of Mt. Fuji (*Fujisan*). This name symbolizes our goal: just as a sea of clouds extends far beyond the mountain peak, UNKAI is designed to be applicable to a more diverse range of proteins than our previous model, FUJISAN.

### 2.4 Performance evaluation

We used several standard metrics to assess the model’s performance: accuracy (*ACC*), precision (*PRE*), recall (*REC*), false positive rate (*FPR*), and F1-score (*F*1). These metrics are defined based on the confusion matrix elements as follows:

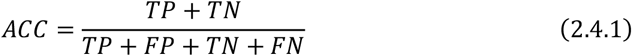

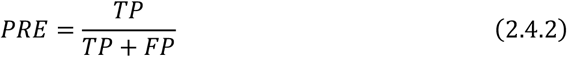

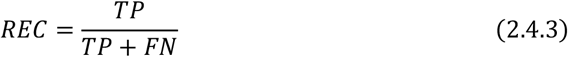

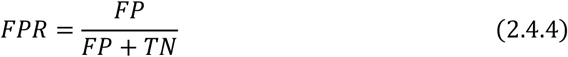

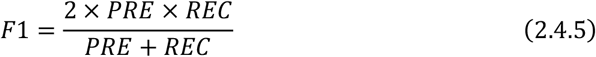

where *TP, FP, TN*, and *FN* represent the numbers of true positives, false positives, true negatives, and false negatives, respectively. Additionally, we utilized the area under the receiver operating characteristic curve (AUROC) and the area under the precision-recall curve (AUPR) to benchmark our model against other existing methods.

### 2.5 Evaluation of the correlation between attention weights and functional annotations

To investigate whether the attention mechanism captures the functional residues of a protein, we extracted experimentally validated functional annotations—including “Active site,” “Binding site,” “Modified residue,” “Natural variant,” and “Site”—for the 59,989 proteins in the test set that carry at least one such annotation from the UniProt database [2]. Let *F*_*A*_ denote the set of residue indices associated with a specific functional annotation *A* within a protein sequence. To perform the residue-level binary classification analysis, we assigned a binary label *y*_*A,j*_ to each residue *j* as follows:

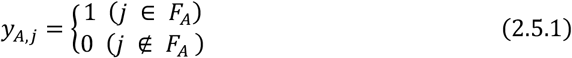

By treating attention weights *α*_*j*_ as classification scores, residue *j* is classified as functional for annotation *A* when *α*_*j*_ is above a threshold. The efficacy of this approach was independently quantified for each annotation *A* via ROC analysis to assess the discriminative power of the attention mechanism across different functional categories.

Since the attention weight profiles exhibited distinct peaks along the protein sequences (see Fig. 3A), we analyzed the positional proximity between these peaks and the known functional residues. Let *I*_top_ denote the set of residue indices with the highest 1% of attention weights within each protein sequence. As a baseline for comparison, we generated a control set *I*_rand_ by randomly sampling indices from the entire sequence without replacement, ensuring it contained the same number of residues as *I*_top_. The minimum sequence distance from residue *j* ∈ *I*_*X*_ (*X* ∈ {top, rand}) to the functional residues in *F*_*A*_ was defined as follows [18]:

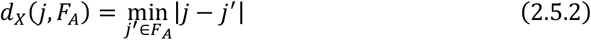

We then calculated the probability mass function *P*_*X,A*_(*k*), representing the probability that *d*_*X*_(*j, F*_*A*_) = *k* across all protein sequences in the test set.

### 2.6 Evaluation of the correlation between attention weights and evolutionary conservation

We also examined the correlations between the attention weights and evolutionary conservation for the proteins in the test set. The evolutionary conservation of residue *j* was quantified using Shannon entropy *H*(*j*), calculated from a multiple sequence alignment (MSA) generated by MMseqs2 [19] as follows:

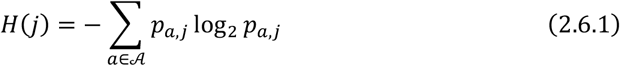

where 𝒜 denotes the set of the 20 standard amino acids, and *p*_*a,j*_ represents the frequency of occurrence of amino acid *a* in the column corresponding to residue *j* . Insertions (represented as lowercase in the A3M format) and gaps (“-” and “.”) were excluded from the entropy calculation. The correlation between the attention weights and the entropies was evaluated via Spearman’s rank correlation coefficient *ρ* [20], which is defined as:

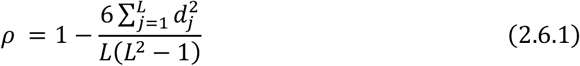

where *L* is the sequence length, and *d*_*j*_ denotes the difference between the ranks of the attention weight and the entropy for residue *j*.

## 3. Results and discussion

### 3.1 Predictive performance of UNKAI

In this study, we developed a deep-learning-based model, UNKAI, designed to predict the functional identity of protein pairs—specifically, whether the two proteins catalyze the same enzymatic reaction involving the same substrate. To construct the dataset, an all-against-all BLAST search was performed on Swiss-Prot sequences associated with Rhea IDs, which provide unique identifiers for enzymatic reactions. From the pairs exhibiting detectable sequence similarity (E-value < 10), we randomly selected 100,000 pairs with the same Rhea ID and 100,000 pairs with different Rhea IDs. The resulting dataset was partitioned into the training, validation, and test subsets at a ratio of 70:15:15. The model was trained for ten epochs, during which stable convergence was observed (Fig. S1). We then assessed its predictive performance and benchmarked it against our previous model, FUJISAN, and other existing methods.

Figure 2 presents a comparison of the ROC and PR curves for these models. Detailed performance metrics, including AUROC and AUPR values, are summarized in Table 1. The results for FUJISAN, Deep-FRI, and the E-value-based baseline were obtained from our previous study [8]. The ROC and the PR curves of UNKAI consistently surpassed those of existing methods, including FUJISAN, achieving AUROC and AUPR values of 0.9930 and 0.9931, respectively. Furthermore, UNKAI exhibited the best performance across all evaluation metrics among the compared methods. These results clearly demonstrate that UNKAI significantly outperformed the existing models in functional identity prediction.

**Figure 2.**
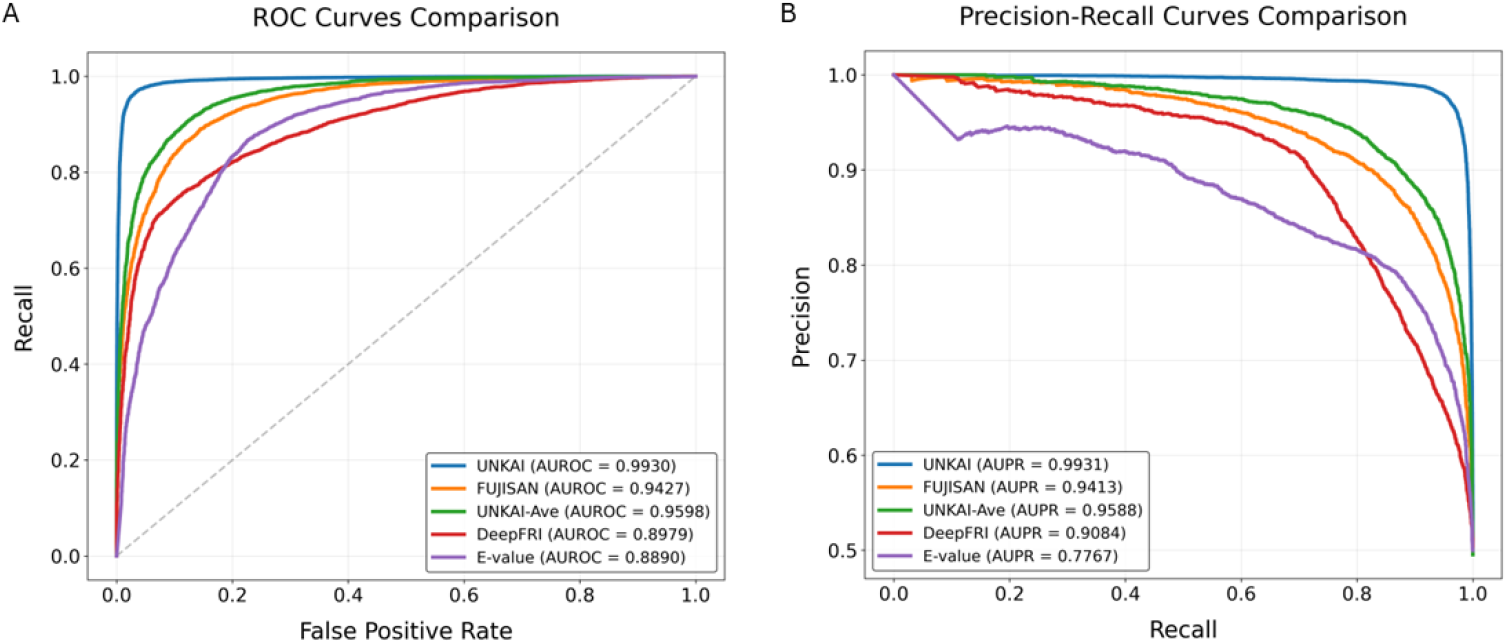
Comparison of predictive performance. (A) Receiver operating characteristic (ROC) curves and (B) precision-recall (PR) curves for UNKAI and benchmark methods. AUROC and AUPR values for each method are provided in parentheses within the legend. The dashed diagonal line in (A) represents the performance of a random classifier.

**Figure 3.**
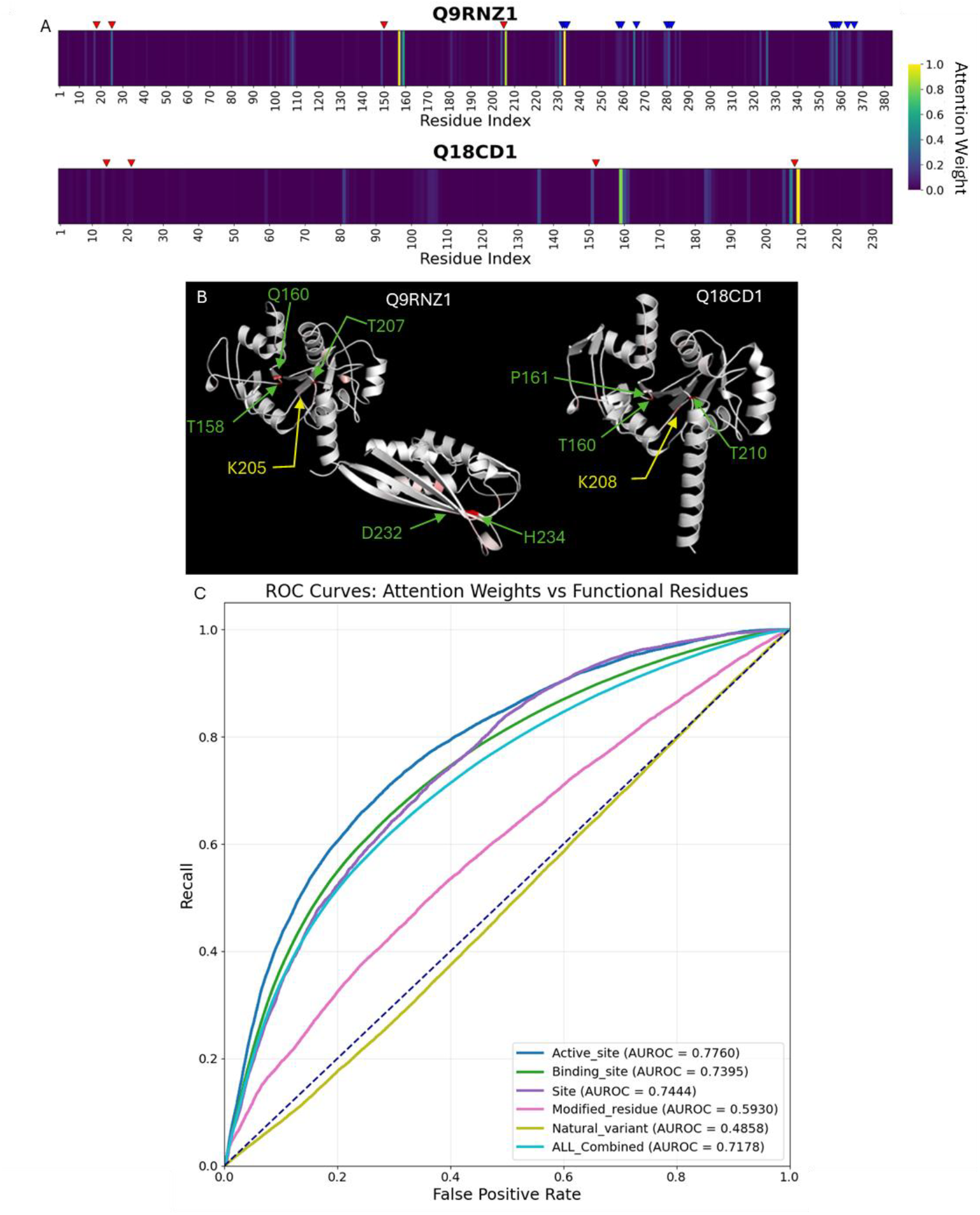
(A) Attention weight heatmaps for proteins Q9RNZ1 and Q18CD1. Functional annotations are indicated by red (Site) and bule (Binding site) triangles above the heatmaps. AlphaFold-predicted structures of Q9RNZ1 and Q18CD1, with residues colored on a gradient from white (low) to red (high) based on their attention weights. Key residues with high attention weights are labeled and indicated with arrows. (C) Receiver operating characteristic (ROC) curves for the prediction of specific functional annotations based on attention weights. AUROC values are provided in parentheses within the legend. The dashed diagonal line represents the performance of a random classifier.

**Table 1.** Comparison of predictive performance between UNKAI and existing methods.

| Method | ACC | PRE | REC | FPR | F1 | AUROC | AUPR |
| --- | --- | --- | --- | --- | --- | --- | --- |
| UNKAI | **0.9666** | **0.9687** | **0.9643** | **0.03110** | **0.9665** | **0.9930** | **0.9931** |
| UNKAI-Ave | 0.8916 | 0.8651 | 0.9279 | 0.1447 | 0.8954 | 0.9598 | 0.9588 |
| FUJISAN<sup>a</sup> | 0.8705 | 0.8724 | 0.8705 | 0.1335 | 0.8728 | 0.9427 | 0.9413 |
| DeepFRI<sup>a</sup> | 0.8088 | 0.7977 | 0.8316 | 0.2145 | 0.8143 | 0.8979 | 0.9084 |
| E-value<sup>a</sup> | 0.8191 | 0.7858 | 0.8775 | 0.2392 | 0.8291 | 0.8890 | 0.7767 |

<sup>a</sup>Cited from Ref. [8].

To evaluate the efficacy of the attention mechanism incorporated into UNKAI, we constructed a variant model, designated as UNKAI-Ave, in which average pooling was employed instead of attention pooling. As shown in Figure 2 and Table 1, the performance of UNKAI-Ave was significantly degraded by the replacement of the attention mechanism, yielding AUROC and AUPR values of 0.9598 and 0.9588, respectively. These scores slightly exceed those of FUJISAN (AUROC: 0.9427, AUPR: 0.9413), confirming that using ESM-C latent representations as input for an MLP-based binary classifier is sufficient to achieve high baseline performance. However, these results highlight that the attention mechanism makes a significant contribution to further enhancing prediction accuracy. In the next section, we investigate how the attention mechanism captures functionally important regions within a protein sequence.

### 3.2 Correlation between attention weights and functional annotations

In the attention layer of UNKAI, a weighted sum of residue embeddings over the sequence length is computed to obtain a fixed-length aggregated feature vector [see Eq. (2.2.3)]. This architecture suggests that residues critical for functional identity are likely to be assigned larger attention weights. For example, UNKAI correctly predicted the functional identity of the protein pair Q9RNZ1 and Q18CD1, which share the same Rhea ID (13429). Notably, residues T158, Q160, K205, T207, D232, and H234 of Q9RNZ1, as well as T160, P161, K208, and T210 of Q18CD1, exhibited high attention weights (Fig. 3A). In the AlphaFold-predicted structures, the highlighted residues of Q18CD1 (T160, P161, K208, and T210) and a subset of those in Q9RNZ1 (T158, Q160, K205, and T207) are spatially situated within the corresponding cavities of each protein (Fig. 3B). Among these, K205 (Q9RNZ1) and K208 (Q18CD1) are specifically annotated as “Site” in UniProt, with the functional description: “position for the nucleophilic attack of MEP.” Interestingly, D232 and H234 of Q9RNZ1, which also exhibit high attention weights, are located in a separate domain. They are annotated as “Binding sites” for a different reaction (Rhea ID: 23864), suggesting that this additional domain catalyzes a reaction specific to Q9RNZ1. Based on these observations, we systematically assessed whether known functional sites could be predicted using the attention weights assigned to the proteins in the test set.

Figure 3C presents the ROC curves for the prediction of functional annotations: “Active site,” “Binding site,” “Modified residue,” “Natural variant,” and “Site.” The attention weights effectively captured “Active site,” “Binding site,” and “Site,” achieving AUROC values of 0.7760, 0.7395, and 0.7444, respectively. In contrast, the predictive performance was notably lower for “Modified residue” (AUROC: 0.5930) and “Natural variant” (AUROC: 0.4858). These results indicate that while the attention weights capture functional sites reasonably well, they do so with varying degrees of efficacy across categories. This further suggests that “Modified residue” and “Natural variant” may be less critical for the determination of functional identity.

The heatmaps (Fig. 3A) reveal that the attention weight profiles exhibit distinct peaks along the protein sequences. This suggests a positional proximity between these peaks and the functional sites, although they do not always coincide exactly. Therefore, we next analyzed the sequence distances between the residues with the highest attention weights and their closest functional sites. Specifically, we compared the distance distributions for the residues in the top 1% of attention weights (*I*_top_) against those for residues randomly selected from the sequences (*I*_rand_) in the test set. For the “Active site,” “Binding site,” and “Site” annotations, the distances between the *I*_top_ residues and their closest functional sites were significantly shorter than those calculated for *I*_rand_ (Fig. 4). Notably, the probability of a residue annotated as “Active site,” “Binding site,” or “Site” existing within five residues of an *I*_top_ residue was 45.9%, whereas the corresponding probability for *I*_rand_ was only 14.4%. For the “Modified residue” and “Natural variant” annotations, however, the distance distributions for *I*_top_ were nearly indistinguishable from those of *I*_rand_, indicating that the attention weights show little to no correlation with these specific annotations. Overall, these findings demonstrate that the attention weights effectively capture functionally important regions within protein sequences.

**Figure 4.**
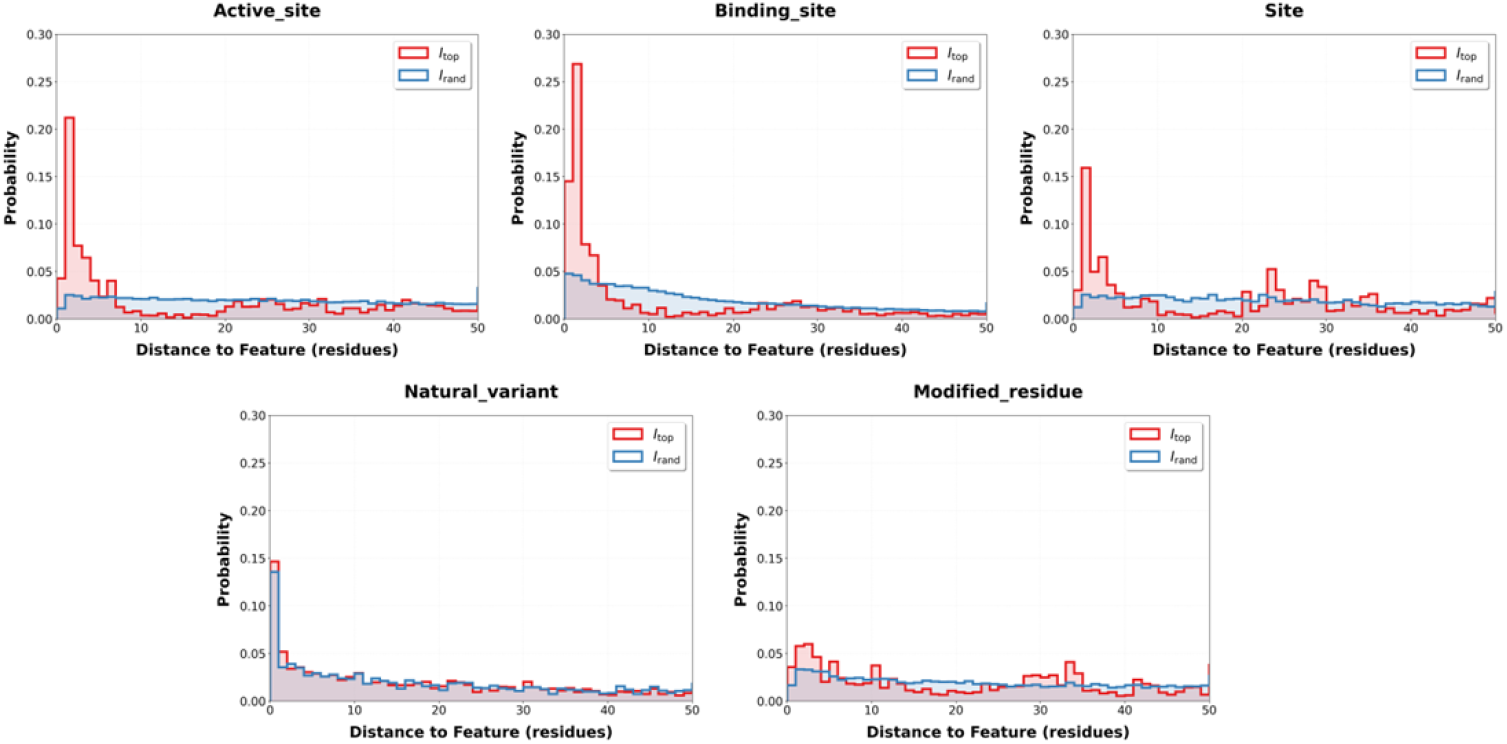
Probability mass functions of the distance from residues to functional sites. The plots show the probability mass functions of the distance from residues with the top 1% attention weights (*I*_top_ red) or randomly selected residues (*I*_rand_ blue) to residues with specific functional annotations. These functions were calculated across all proteins with functional annotations in the test set.

### 3.3 Correlation between attention weights and evolutionary conservation

Since functionally important residues tend to be evolutionarily conserved [21], we assessed the correlation between attention weights and evolutionary conservation using Spearman’s rank correlation coefficients. Evolutionary conservation was quantified using Shannon entropy calculated from the multiple sequence alignment obtained for each protein in the test set. Figure 5A presents a scatter plot of attention weights against entropy for all residues. As discussed in the previous section, higher attention weights reflect functional importance, whereas lower entropy values indicate higher degrees of conservation. Consistently, the plot indicates a negative correlation. The Spearman’s rank correlation coefficients calculated for each protein exhibited a unimodal distribution with a mean value of −0.4421 (Fig. 5B), indicating that attention weights are moderately correlated with evolutionary conservation. Notably, this result suggests that evolutionary conservation alone is not sufficient for accurate functional identity prediction. Considering that the correlations between attention weights and functional annotations were also moderate, as observed in the previous section, it is evident that the attention mechanism integrates information extending far beyond simple evolutionary conservation and functional annotations through the learning process.

**Figure 5.**
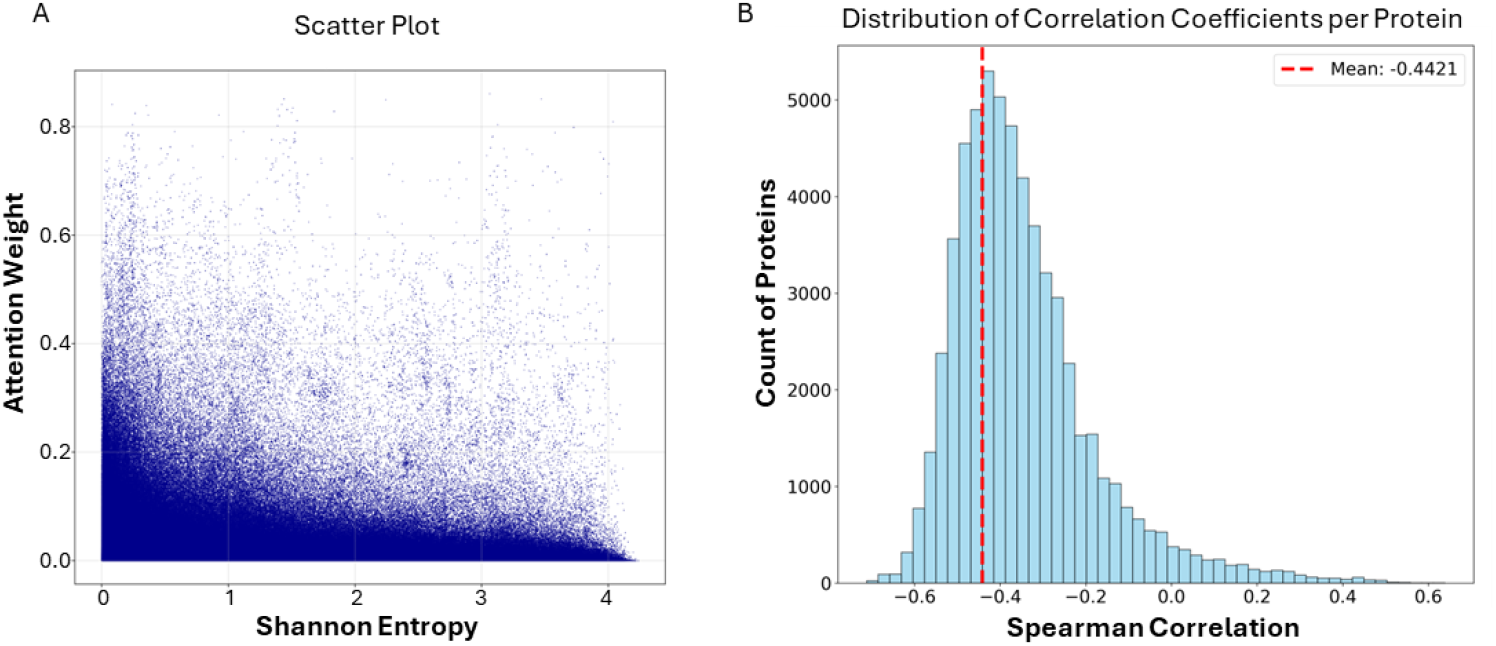
Correlation between attention weights and evolutionary conservation. (A) Scatter plot of per-residue attention weights against Shannon entropies across all proteins in the test dataset. Lower Shannon entropy values indicate greater conservation. (B) Distribution of Spearman’s rank correlation coefficient between Shannon entropies and attention weights calculated for each protein. The dashed red line indicates the mean correlation coefficient of −0.4421.

### 3.4 Performance on low sequence similarity dataset

Since UNKAI and FUJISAN were trained on datasets consisting of protein pairs with detectable sequence similarity (E-value < 10), their predictive performance may inherently depend on the degree of sequence similarity between the input pairs. In our previous study, we demonstrated the robustness of FUJISAN by evaluating its performance on protein pairs with low full-length similarity [8]. To evaluate UNKAI’s performance in this context, we constructed a low sequence similarity (LSS) dataset comprising pairs with E-values > 10^−10^ (680 pairs with identical functions and 680 pairs with different functions). To ensure a rigorous and independent evaluation, we confirmed that this LSS dataset contains no overlap with the training, validation, or test sets used in the original experiments. Figures 6A and 6B present the ROC and PR curves of UNKAI, UNKAI-Ave, FUJISAN, and other existing methods. Although the AUROC and AUPR values of UNKAI (0.9673 and 0.9749, respectively) were slightly lower than those obtained on the original test set, UNKAI still outperformed all other models on the LSS dataset. These results suggest that UNKAI captures essential functional features that extend beyond simple sequence homology. While alignment-based methods, such as those relying on E-values, are inherently limited by sequence similarity, our model demonstrates high robustness in identifying functional relationships even for “remote homologs” with low sequence similarity.

**Figure 6.**
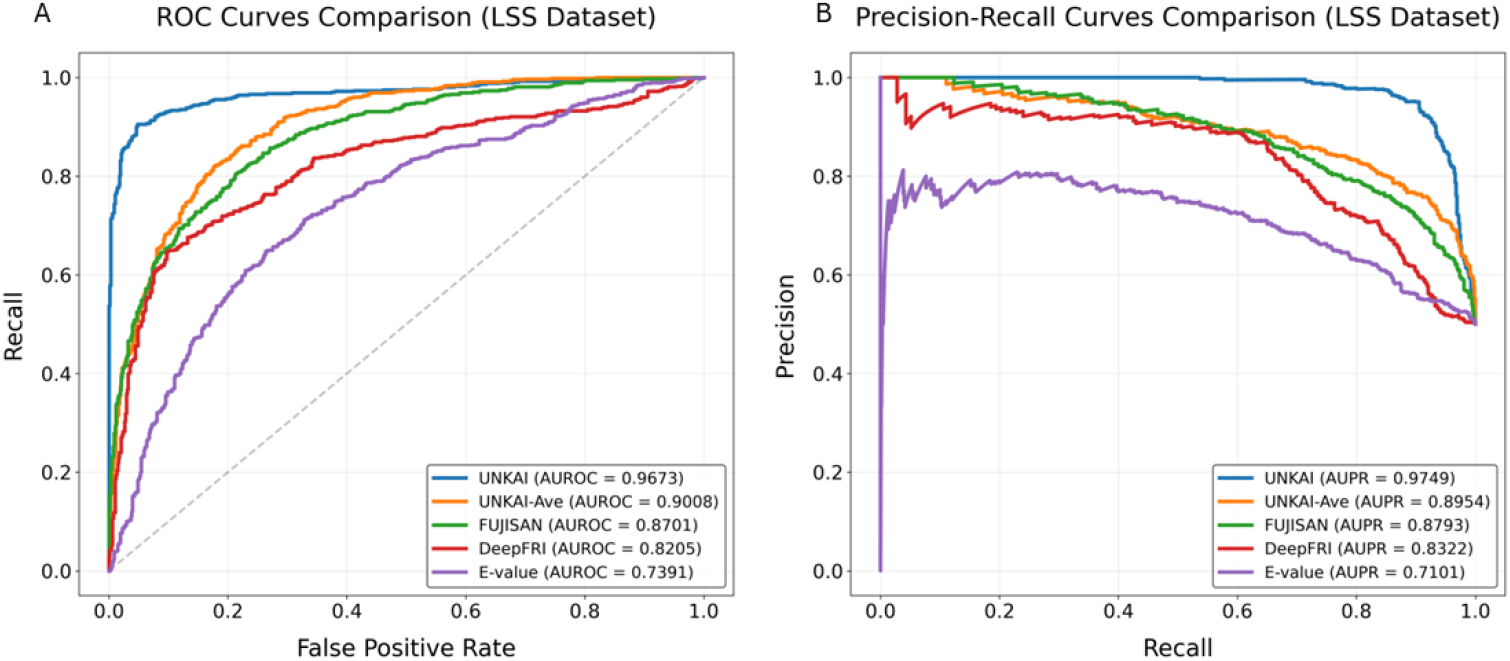
Predictive performance comparison on the low sequence similarity (LSS) dataset. (A) Receiver operating characteristic (ROC) curves and (B) precision-recall (PR) curves for UNKAI and benchmark methods evaluated on the LSS dataset. AUROC and AUPR values for each method are provided in parentheses within the legend. Results of FUJISAN, DeepFRI, and E-value are cited from Ref. [8]. The dashed diagonal line in (A) represents the performance of a random classifier.

### 3.5 Limitations

In general, prediction accuracy is inherently limited for inputs that lack similar samples in the training set. Although the training set of UNKAI encompasses a wide variety of proteins, the distribution of functional categories—classified at the third digit level of Enzyme Commission (EC) numbers—exhibits some bias, reflecting the distribution of the original database (Fig. S2). To evaluate UNKAI’s predictive capabilities for entirely unseen EC subclasses, we conducted a leave-one-subclass-out evaluation for five specific subclasses (Table 2). Notably, predictive performance for these unseen subclasses was lower than the overall performance reported in Table 1. If UNKAI relied solely on broad sequence homology, it would maintain consistent performance even for unknown categories, provided that some sequence similarity exists. However, the observed decline in performance suggests that UNKAI captures essential functional features specific to each category, extending beyond simple sequence homology. This result further underscores that the attention mechanism integrates complex, category-specific information through the learning process, which in turn necessitates a large, functionally diverse training set to achieve broad generalization.

**Table 2.** Predictive performance of UNKAI on unseen EC subclasses.

| **Model<sup>a</sup>** | **ACC** | **PRE** | **REC** | **F1** | **AUROC** | **AUPR** |
| --- | --- | --- | --- | --- | --- | --- |
| UNKAI-EC3.6.4 | 0.7629 | 0.8301 | 0.6610 | 0.7360 | 0.8489 | 0.8542 |
| UNKAI-EC2.7.1 | 0.7388 | 0.8255 | 0.6055 | 0.6986 | 0.8312 | 0.8340 |
| UNKAI-EC2.7.4 | 0.6655 | 0.6520 | 0.7102 | 0.6798 | 0.7337 | 0.7302 |
| UNKAI-EC3.6.1 | 0.7000 | 0.7560 | 0.5906 | 0.6631 | 0.7614 | 0.7686 |
| UNKAI-EC4.2.1 | 0.7553 | 0.7378 | 0.7921 | 0.7640 | 0.8464 | 0.8719 |

<sup>a</sup>The suffix “-ECX.Y.Z” denotes the specific EC subclass that was excluded from the training set and subsequently used for performance evaluation.

## 4. Conclusions

In this study, we developed UNKAI, a deep-learning-based model designed to predict the functional identity of protein pairs by leveraging ESM-C latent representations of the protein sequences and an attention mechanism. Unlike our previous hypothesis-driven method, FUJISAN, UNKAI adopts a fully data-driven approach to dynamically capture residue-level importance. Our results demonstrate that UNKAI significantly outperforms existing methods, including FUJISAN. This superior performance is largely attributed to the attention mechanism, which computes weighted sums of the ESM-C latent representations of input protein sequences to yield fixed-length aggregated feature vectors. Notably, the attention weights exhibited moderate but significant correlations with both evolutionary conservation and functional annotations—such as “Active site,” “Binding site,” and “Site” from the SwissProt database. These findings indicate that while the attention mechanism effectively captures known functional regions, it also integrates additional information that extends far beyond simple conservation patterns through the learning process. Although challenges remain in generalizing to completely unseen functional classes, this framework provides a powerful tool for the functional annotation of uncharacterized proteins and serves as a foundational step toward more generalized enzyme discovery methods.

## CRediT authorship contribution statement

**Kotaro Ukai**: Conceptualization, Methodology, Software, Validation, Formal analysis, Investigation, Data curation, Visualization, Writing - original draft. **Suguru Fujita**: Conceptualization, Methodology, Formal analysis, Investigation, Data curation, Writing - original draft, Writing – review & editing. **Tohru Terada**: Conceptualization, Methodology, Formal analysis, Investigation, Resources, Writing – review & editing, Supervision, Project administration, Funding acquisition.

## Acknowledgments

This work was partially supported by the JSPS KAKENHI Grant Number JP22H05126 and by Research Support Project for Life Science and Drug Discovery (Basis for Supporting Innovative Drug Discovery and Life Science Research (BINDS)) from Japan Agency for Medical Research and Development (AMED) under Grant Number JP26ama121027, and grant provided by Uehara Memorial Foundation. The authors would like to thank Gemini (Google) for its assistance in refining the manuscript’s English phrasing and grammatical structure.

