## Supplementary material for "UNKAI: A protein functional identity prediction model based on ESM-C latent representations and the attention mechanism": Fig. S1, Fig. S2

### Supplementary Figures

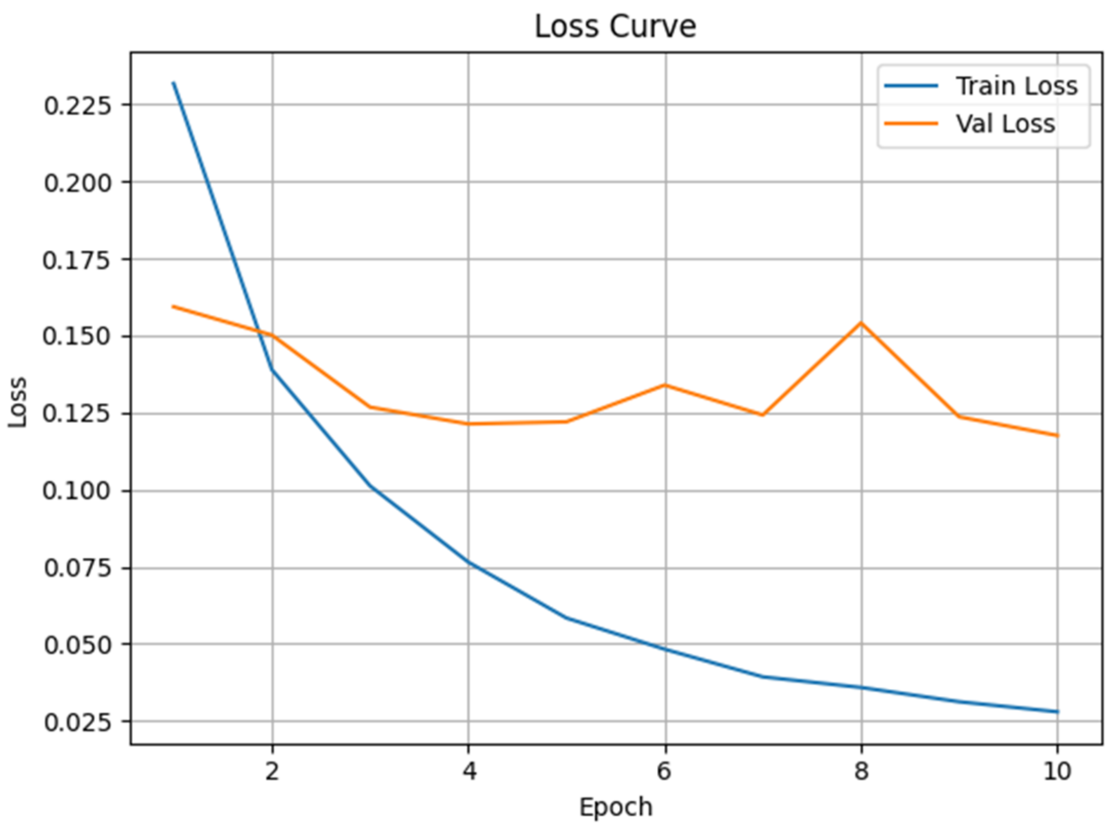


**Figure S1.** Learning curves of the proposed model. The plot shows the training and validation loss per epoch for UNKAI. The binary cross-entropy loss was minimized using the Adam optimizer with a learning rate of $3.33 \times 10^4$.

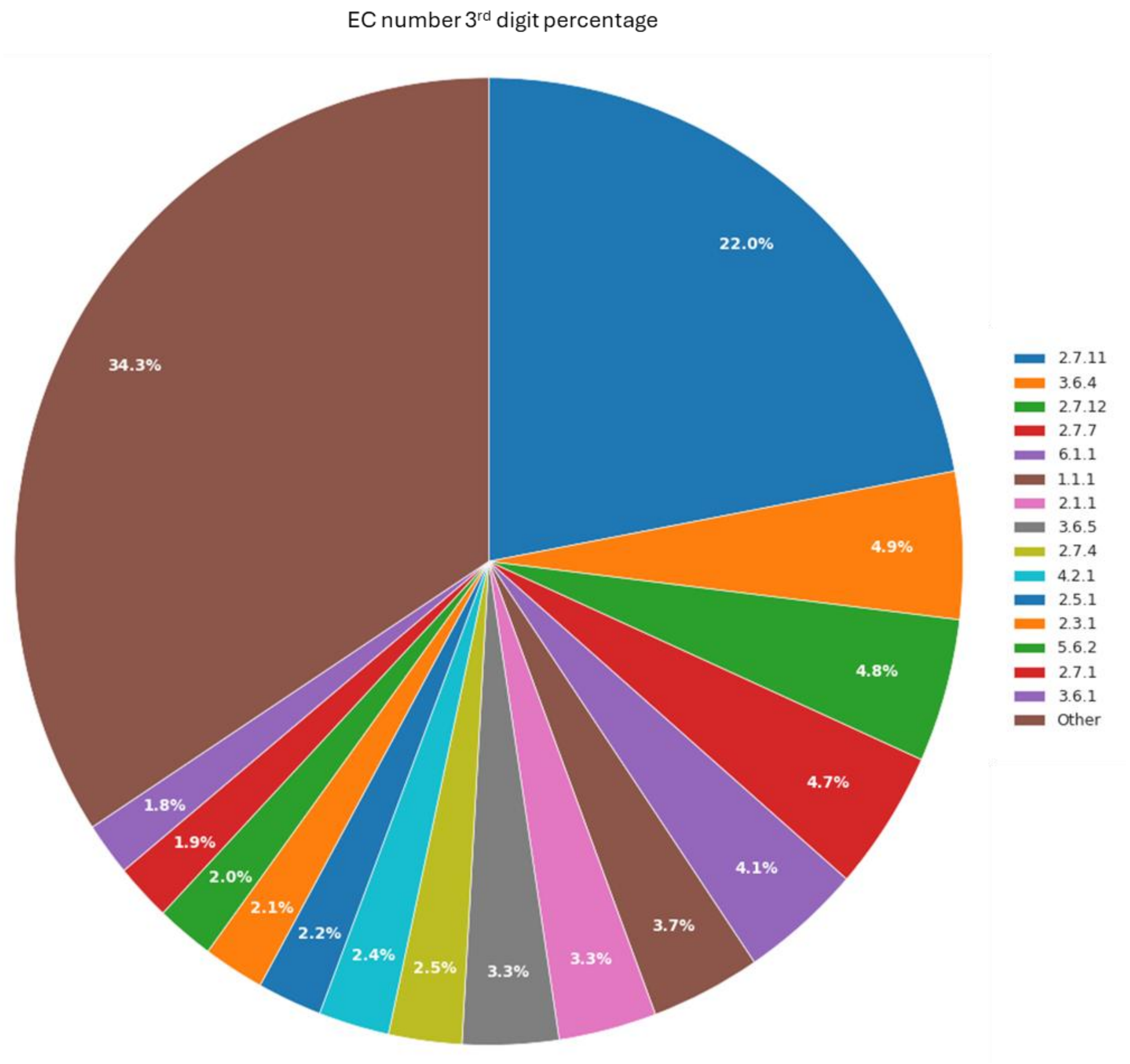


**Figure S2.** Distribution of functional categories in the training set. The pie chart illustrates the proportions of proteins classified by the first three digits of their Enzyme Commission (EC) numbers. Only the top 15 functional categories are explicitly labeled, with the remaining categories grouped under "Other."
